# Inferring the collective dynamics of neuronal populations from single-trial spike trains using mechanistic models

**DOI:** 10.1101/671909

**Authors:** Christian Donner, Manfred Opper, Josef Ladenbauer

## Abstract

Multi-neuronal spike-train data recorded in vivo often exhibit rich dynamics as well as considerable variability across cells and repetitions of identical experimental conditions (trials). Efforts to characterize and predict the population dynamics and the contributions of individual neurons require model-based tools. Abstract statistical models allow for principled parameter estimation and model selection, but possess only limited interpretive power because they typically do not incorporate prior biophysical constraints. Here we present a statistically principled approach based on a population of doubly-stochastic integrate-and-fire neurons, taking into account basic biophysics. This model class comprises an idealized description for the dynamics of the neuronal membrane voltage in response to fast independent and slower shared input fluctuations. To efficiently estimate the model parameters and compare different model variants we compute the likelihood of observed single-trail spike trains by leveraging analytical methods for spiking neuron models combined with inference techniques for hidden Markov models. This allows us to reconstruct the shared input variations, classify their dynamics, obtain precise spike rate estimates, and quantify how individual neurons couple to the low-dimensional overall population dynamics, all from a single trial. Extensive evaluations based on simulated data show that our method correctly identifies the dynamics of the shared input process and accurately estimates the model parameters. Validations on ground truth recordings of neurons in vitro demonstrate that our approach successfully reconstructs the dynamics of hidden inputs and yields improved fits compared to a typical phenomenological model. Finally, we apply the method to a neuronal population recorded in vivo, for which we assess the contributions of individual neurons to the overall spiking dynamics. Altogether, our work provides statistical inference tools for a class of reasonably constrained, mechanistic models and demonstrates the benefits of this approach to analyze measured spike train data.

## Introduction

Cortical computations are represented in the collective spiking activity of multiple neurons. The growing interest to uncover how these neuronal populations process and transform complex incoming information into decisions and motor actions has brought about cell-resolving activity measurements at an increasing scale and pace. Although these activity patterns can, in principle, span a high-dimensional space, often a large fraction of neural variability is captured by low-dimensional manifolds [1–5].

Statistical inference based on generative models is a powerful approach to interpret the measured spike train data and characterize the hidden, low-dimensional, collective dynamics [6, 7]. For this purpose typically abstract, time-dependent latent variable models are fitted to the data [6, 8–15]. These approaches are well suited for quantifying the structure in the data, and benefit from statistically principled parameter estimation and model selection methods. However, their interpretive power and capacity for identifying neural mechanisms are limited as the underlying models typically do not incorporate prior biophysical constraints.

Mechanistic models of neural populations, on the other hand, involve variables and parameters that can be biophysically interpreted, and have proven useful to dissect neural dynamics. Well-known exponents are the detailed Hodgkin–Huxley-type neuron models [16, 17], which can involve complex morphology and patterns of ionic currents; however, they are not well suited to analyze spike-train data, because multiple combinations of parameters give rise to the same firing patterns [18, 19]. An alternative, prominent class of models with reduced complexity are spiking neuron models of the *integrate-and-fire* (I&F) type, which implement in a simplified manner essential biophysical constraints. These models have been advanced in recent years to accurately predict neuronal activity [20–22] and classify multiple neuron types [17, 23, 24]. They have become state-of-the-art models for describing neural activity in in-vivo-like conditions and have been applied in a multitude of studies on neural network dynamics. Yet, while I&F models are biophysically more faithful than abstract statistical models, fitting such models to multi-neuronal spike trains with account of variability and latent, low-dimensional population dynamics, is a substantial challenge.

Here we consider a population of doubly-stochastic I&F neurons to model highly non-stationary, collective spiking dynamics as typically observed in vivo. The hidden neuronal inputs, which impinge on the hidden membrane potentials, contain fast independent fluctuations that give rise to spiking variability, and slower shared variations that dominate the low-dimensional, latent population dynamics. Shared variability across observed neurons, which typically reflect only a small fraction of the local population, is captured in the model by a common, latent process rather than by putative direct coupling. We present a statistically principled approach based on derived likelihood functions to fit this type of model to single-trial spike trains and quantitatively compare different model variants, including simpler phenomenological models. Specifically, we efficiently compute the likelihood of a given spike train by exploiting analytical methods for stochastic I&F neuron models [25] combined with inference techniques for hidden Markov models [26]. This allows us to infer the model parameters by likelihood maximization, classify the latent dynamics via likelihood-based model selection, and estimate hidden time series from the time-varying probability density over the latent state.

We evaluate our approach extensively on synthetic data in terms of reconstruction of the true latent time series, classification of their dynamics, and estimation of the model parameters. We then validate our method using in-vitro ground truth recordings of cortical neurons stimulated by current signals with additive noise [27]. We successfully reconstruct the true dynamics of the signal and demonstrate improved fitting performance compared to a classical inhomogeneous Poisson point process model. Finally, we apply our approach to extracellular recordings from macaque primary visual cortex in vivo [28] to characterize the low-dimensional population dynamics and the contributions of individual neurons.

## Results

Our results are structured as follows. The model is explained in section 1. In the following two sections we outline our inference methods and evaluate them exhaustively on synthetic data. For reasons of clarity and comprehensibility we first focus on single neurons in section 2 and consider neuronal populations in section 3. In section 4 we validate our approach based on in-vitro recordings. In section 5 we apply our methods to population spike train data from extracellular recordings in vivo.

### 1 Statistical modeling with doubly-stochastic I&F neurons

We consider having observed cell-resolved spike trains (ordered sets of spike times) from a population of *N* neurons. Such data are typically obtained from extracellular multi-electrode recordings after pre-processing that includes spike sorting [29]. The activity of each neuron (with index *i*) of the population shall be described by the classical leaky I&F model [30] with membrane time constant *τ*_m_, where the compound synaptic input is given by a Gaussian white noise process with mean

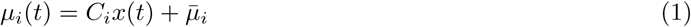

and standard deviation *σ*_*i*_. This process effectively models synaptic bombardment impinging on the neuronal membrane voltage (see, e.g., [31–34]). The input includes shared, slow variations *x*(*t*) which may be caused by a common component in the external drive or by network interactions, and independent, rapid fluctuations with strength *σ*_*i*_. 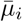 is a cell-specific offset. *C*_*i*_ quantifies the extent to which neuron *i* is affected by *x*(*t*). Notably, in the context of inferring *x*(*t*) from observed activity data, *C*_*i*_ measures how much the activity of neuron *i* contributes to the shared variations. In the following, we simply refer to it as *coupling strength* (for the coupling between the individual activity of a neuron and the shared dynamics).

Since the dynamics of the common input are not known we describe *x*(*t*) by a stochastic process. In particular, we consider two qualitatively different Markov models, an *Ornstein-Uhlenbeck process* (OUP) and a *Markov jump process* (MJP), which gives rise to two model variants. For the OUP *x*(*t*) varies continuously with time constant *τ*, whereas for the MJP *x*(*t*) is piece-wise constant intermittently jumping to different values with rate *τ*^-1^. The stationary probability density of both processes is standard normal and their autocorrelation functions are identical. For further details on the generative model see Methods section 1. In Results sections 4 and 5 below we also consider a simpler, classical model for comparison, which describes the spike train of neuron *i* by a Poisson point process with rate exp(*µ*_*i*_(*t*)), where *µ*_*i*_(*t*) is given by Eq (1).

It is important to point out that not all model parameters need to be estimated. The membrane voltage can be scaled such that the remaining parameters of interest for inference are those for the input together with the membrane time constant (see Methods section 1). Moreover, a change of *τ*_m_ can be well compensated for in terms of spiking dynamics by appropriate changes of 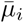 and *σ*_*i*_ [25]. Therefore, we focus on the following parameters for inference 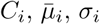 for *i ∈* {1, …, *N*} and *τ*, with particular interest on the coupling strengths (*C*_*i*_) and the time constant (*τ*).

### 2 Inference for single neurons

#### Outline of inference approach

It is instructive to first consider a single neuron. For improved readability we omit the neuron index *i* and group the parameters as 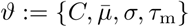, and *τ*. We collect the measured spike-train data in the increasing sequence of spike times *t*_0:*K*_ := (*t*_0_, …, *t*_*K*_) and define the *k*-th interspike interval (ISI) by *s*_*k*_ := *t*_*k*_ - *t*_*k-*1_. *K* is the number of observed ISIs. For notational ease below we use that *s*_*k*_, the duration of the *k*-th ISI, implicitly contains information about the start time and end time of the interval. As spike emission in the leaky I&F model is a renewal process the likelihood of observing a given spike train from the model is completely determined by the likelihoods of observing the constituent ISIs. To tackle the inference problem we approximate the time series of the process *x*(*t*) for each ISI by one value, *x*_*k*_ for ISI *s*_*k*_, which is justified by the assumption that *x*(*t*) varies slowly, i.e., *τ* is large compared to the average ISI. Defining *x*_0_ as the value of the process at *t* = 0 and *x*_*k*_ as the value at *t* = *t*_*k*_ for *k* ≥ 1 we obtain a hidden Markov model with sequence of latent states *x*_0:*K*_ := (*x*_0_, …, *x*_*K*_). The observations are contained in the sequence *s*_1:*K*_ := (*s*_1_, …, *s*_*K*_). The joint likelihood of observing a given spike train and realization of *x*(*t*), approximated by sequence *x*_0:*K*_, is given by

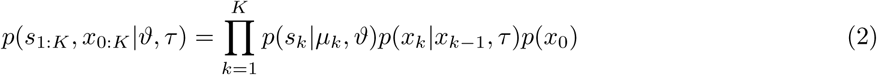

with effective mean input 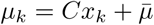. *p*(*s*_*k*_|*µ*_*k*_, *ϑ*) is the ISI probability density of the leaky I&F neuron exposed to Gaussian white noise input (with mean *µ*_*k*_ and standard deviation *σ*), evaluated at ISI *s*_*k*_. This density can be accurately computed by solving a Fokker-Planck partial differential equation, which can be achieved numerically in efficient ways [25]. *p*(*x*_*k*_|*x*_*k-*1_, *τ*) is the transition probability density for *x*(*t*), from state *x*_*k-*1_ to state *x*_*k*_, which depends on the time constant *τ* and on the time duration between states *x*_*k-*1_ and *x*_*k*_. For uncluttered notation this duration is not explicitly indicated. The transition densities for both model variants (OUP and MJP) are known (for details see Methods section 2). *p*(*x*_0_) is the prior probability density of the process at *t* = 0 < *t*_0_ (prior to *t*_0_) for which we assume its stationary distribution, a standard normal. Note that in Eq (2) we have used the renewal property of leaky I&F neurons. We obtain the likelihood of the observations by marginalizing Eq (2) with respect to *x*_0:*K*_,

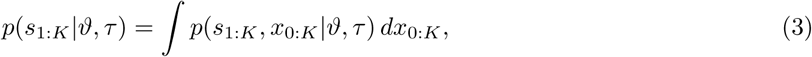

which is efficiently performed by an iterative procedure known as forward filtering (see Methods section 3). Parameter estimates 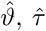 are determined by maximizing the likelihood,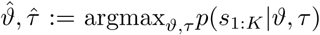. For this purpose it is an important advantage that we can evaluate *p*(*s*_1:*K*_ |*ϑ, τ*) with high accuracy and low computational effort. For maximization we use an established simplex-based method [35] (for details see Methods section 3).

After having inferred the parameter values we compare the fitting performance of both model variants (OUP and MJP) to identify the best model and thereby classify the dynamics of the slow input variations. Specifically, we use the log-likelihood ratio (LLR) for comparison, equivalently expressed as the difference

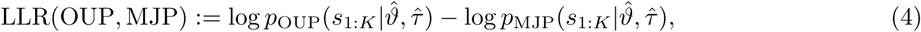

where subscripts OUP and MJP indicate the respective process. Positive values of the LLR indicate that the OUP model variant is favored, while negative values indicate that the MJP model variant better describes the data.

We further compare the doubly-stochastic I&F model to a Poisson point process whose principal parameter, the rate, is given by *λ*(*t*) = exp(*µ*(*t*)) with 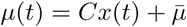 (cf. Eq (1)) and *x*(*t*) is described by an OUP or MJP as specified above. The exponential function is a typical choice for the mapping between input and output rate in Poisson models of this type, see e.g. [10, 14, 36]. We infer the parameters of this model similarly as for the I&F model, the only two differences are that for the Poisson model the ISI density is explicitly expressed by 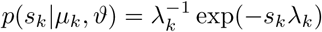 with 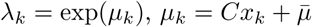, and *ϑ* contains two parameters, *C* and 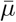, instead of four. For model comparison we use the Akaike information criterion (AIC), which takes into account both goodness of fit and complexity of a model (for details see Methods section 4).

Finally, to reconstruct the time series of the process *x*(*t*) and estimate the instantaneous spike rate of the neuron we infer 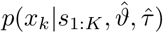, the probability density of the state *x*_*k*_ during the ISI *s*_*k*_ given the observations *s*_1:*K*_ and the inferred parameter values, for each *k*. We compute this density using an iterative technique known as backward smoothing, which in addition to the above-mentioned forward filtering method constitutes the established forward-backward algorithm for hidden Markov models [26] (see Methods section 5). Our estimate for the time series of *x*(*t*) is then obtained by the sequence of expected values (⟨*x*_0_⟩, …, ⟨*x*_*K*_⟩) calculated using these densities. Additionally, we estimate the spike rate time series by the sequence (*r*_0_, …, *r*_*K*_) of expected inverse mean ISIs, where the mean ISI for the *k*-th interval is calculated using *p*(*s*|*µ*_*k*_, *ϑ*) and the expectation is with respect to *p*(*x*_*k*_|*s*_1:*K*_, *ϑ, τ*) (for details see Methods section 5). Note that *r*_*k*_ represents the instantaneous spike rate at the time that corresponds to *x*_*k*_.

#### Evaluation on synthetic data

For evaluation we consider synthetic spike-train data from numerical simulations of the underlying generative model. We assess the performance of the proposed method to reconstruct the time series of *x*(*t*), classify its dynamics and estimate the parameters (Fig 1). Example spike trains and true time series of *x*(*t*) together with the inferred sequences of conditional density over the latent state are shown in Fig 1A,B. The evolution of the density inferred from the spike train well resembles the true time series. Using the incorrect model variant for inference leads to an approximation that appears slightly less accurate.

**Figure 1.**
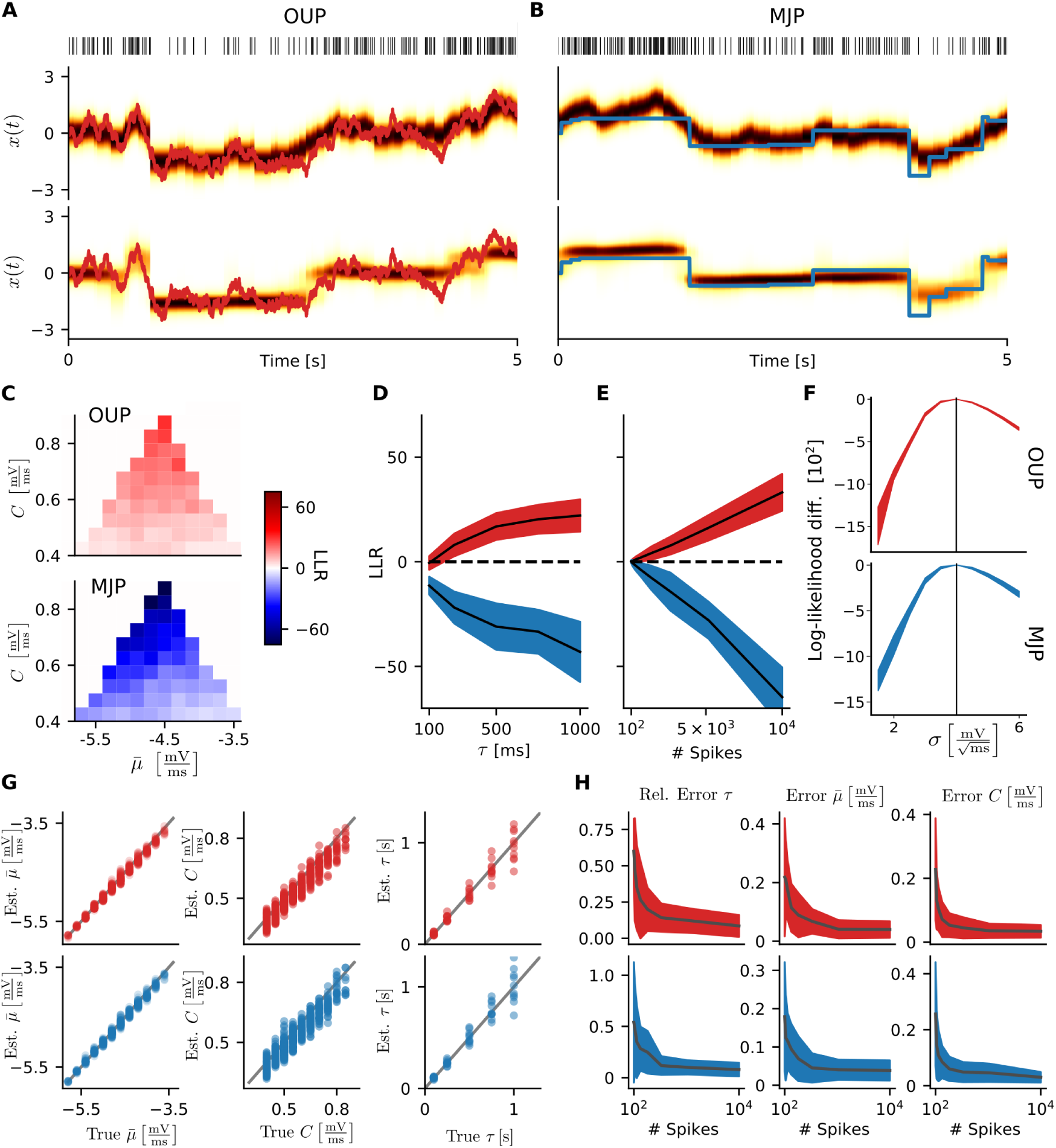
Inference results for single neurons using synthetic data. **A**: Example spike train from the doubly-stochastic I&F model with OUP *x*(*t*) (red traces below) together with sequence of estimated density over the latent variable (color map) when using the OUP (center) or MJP (bottom) for inference. **B**: Example as in **A** with true *x*(*t*) given by a MJP (blue traces). **C**: Log-likelihood ratio (LLR, cf. Eq (4)) for different values of 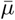 and *C* with true *x*(*t*) given by an OUP (top) or by a MJP (bottom). We considered all 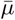, *C*-pairs, for which the expected spike rate remains in the interval [1, 110] Hz for at least 98 % of the simulation time. This was analytically determined from the steady-state spike rate and the quantiles of the stationary distribution of *µ*(*t*). Results represent averages over 10 realizations. In **D**–**H** red color indicates that the true *x*(*t*) is given by an OUP, blue color indicates that it is a MJP. For each parametrization results from 10 realizations are shown, colored areas denote mean ± standard deviation. **D**: LLR as a function of time constant *τ*. **E**: LLR as a function of number of observed spikes. **F**: Difference between maximal log-likelihood for different values of *σ* and the one obtained for the true value. **G**: Estimated versus true parameter values of 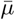 (left), *C* (center) and *τ* (right). The results for 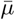 and *C* correspond to the fits in **C**, the results for *τ* correspond to the fits in **D**. **H**: Relative and absolute errors between estimated and true parameter values as a function of number of observed spikes, respectively. The results correspond to the fits in **E**. If not stated otherwise, parameter values for generating the data were 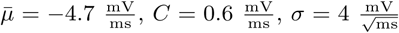 and *τ=* 500ms 5 × 10^3^ observed spikes were used for fitting.

A systematic quantification of classification accuracy for a wide range of parameter values and amount of observed data is presented in Fig 1C-E. Classification according to the LLR is correct throughout the tests. The decision becomes more obvious as the coupling strength *C*, the time constant *τ* or the length of the spike train increases. Intuitively, it is challenging to classify the dynamics if the neuron is only weakly affected by *x*(*t*) (small *C*) or if *x*(*t*) varies rapidly (small *τ*). Surprisingly, a few hundred observed spikes are already sufficient to identify the latent process. Note the small bias towards the MJP, which may be caused by the piece-wise constant approximation of the process in our inference method.

Accuracy of parameter inference is visualized in Fig 1F-H. The true values of the parameters 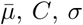, and *τ* are well recovered, and estimation errors decrease with increasing spike train length. Note that the membrane time constant *τ*_m_ was not estimated here, but set to the true value. Setting *τ*_m_ instead to a wrong value within a biologically plausible range affects the maximal likelihood only very little (see Fig S1), because a change of *τ*_m_ can be well compensated for in terms of spiking dynamics by suitable changes of the input parameters (cf. [25]). We, therefore, keep *τ*_m_ fixed for the rest of this study.

In sum, our method allows for accurate parameter inference and characterization of the latent dynamics based on synthetic data of single neurons. Even relatively short spike trains lead to satisfactory results.

### 3 Inference for neuronal populations

#### Outline of inference approach

We now turn to a larger population of neurons as described in Results section 1. We collect the observed data in terms of (cell-resolved) ISIs of all *N* neurons in the single sequence *s*_1:*K*_ := (*s*_1_, …, *s*_*K*_) which is ordered according to the last spike time of each ISI. Similarly as for single neurons we effectively approximate the time series of *x*(*t*) for each neuron and each ISI by one value, which is justified by the slowness of the process. Specifically, we define the sequence of latent states *x*_0:*K*_ := (*x*_0_, …, *x*_*K*_) where *x*_0_ = *x*(0) and *x*_*k*_ = *x*(*t*_*k*_) is the value of the process at the *k*-th spike time across the population for *k* ≥1. Let *i*_*k*_ indicate the neuron that corresponds to the *k*-th ISI *s*_*k*_ in the sequence *s*_1:*K*_, ending at spike time *t*_*k*_. The mean input for neuron *i*_*k*_ across ISI *s*_*k*_ is thus approximated by 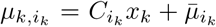. That is, each neuron “samples” the latent state sequence *x*_0:*K*_ (which can contain multiple values within an ISI) in a distinct way according to its spike times. This approximation allows us to extend the approach for a single neuron (cf. Results section 2) to a neuronal population in a straightforward way. We express the joint likelihood of the observed data and the latent state sequence for this hidden Markov model as

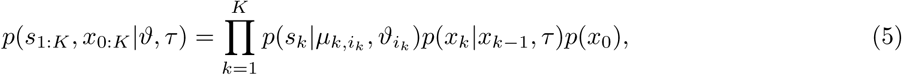

where 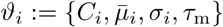 summarizes neuron-specific parameters, *ϑ* := {*ϑ*_1_, …, *ϑ*_*N*_}. Note that the three probability density functions on the right hand side of Eq (5) are identical to those of Eq (2) explained in Results section 2. To obtain the likelihood of observing the data from the model we marginalize Eq (5) with respect to *x*_0:*K*_, and then continue analogously to the single neuron case for inference of model parameters, classification of the dynamics of *x*(*t*), reconstruction of its time series and estimation of the time-varying population spike rate (for details see Methods sections 3–5). Notably, although we can accurately and rapidly evaluate the (marginalized) likelihood using numerical methods (cf. Methods sections 2 and 3), maximization with respect to 3*N* + 1 parameters becomes a serious challenge for a large population. Therefore, we use an efficient, approximate technique for optimization (see Methods section 3).

#### Evaluation on synthetic data

We now evaluate our method using simulated data from the doubly-stochastic I&F population model (Fig 2). Example population spike trains, spike rate histograms, hidden time series of *x*(*t*), and the results from our method are shown in Fig 2A-D. In all cases the inferred sequences of conditional density over the latent state capture well the true time series and the population rate estimates match with the empirical rate histograms.

**Figure 2.**
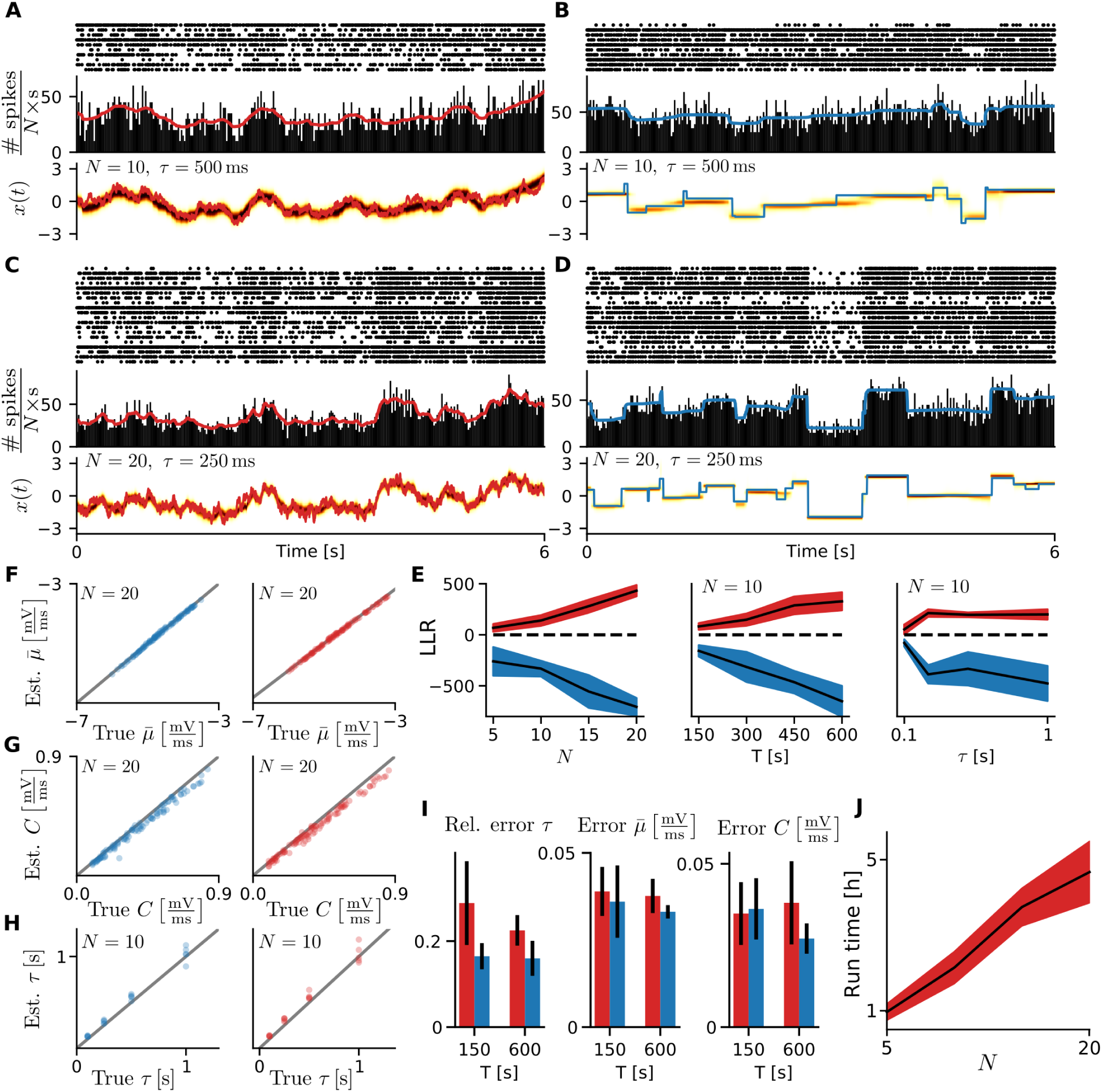
Inference results for neuronal populations using synthetic data. **A**, top: Spike trains from the doubly-stochastic I&F population model with OUP *x*(*t*) for *N* = 10 and *τ* = 500 ms (6 s shown). Center: Population spike rate histogram and sequence of estimated instantaneous population spike rate (red trace). Bottom: True time series of *x*(*t*) (red trace) and sequence of the estimated density over this latent variable (color map). **B**: Example as in **A** with true *x*(*t*) given by a MJP (blue trace, bottom). **C** and **D**: Examples as in **A** and **B**, respectively, for *N* = 20 and *τ* = 250 ms. In **E**–**J** red color indicates that the true *x*(*t*) is given by an OUP, blue color indicates that it is a MJP. For each parametrization results from 5 realizations are shown. **E**: LLR as a function of number of neurons *N* (left), recording duration *T* (center), and time constant *τ* (right). Colored areas denote mean *±* standard deviation. **F**–**H**: Estimated versus true parameter values of 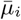 and *Ci* for *i* ∈ {1, …, *N*}, and *τ*. The results for *τ* correspond to the fits in **E**, right. **I**: Relative and absolute errors between estimated and true parameter values for two recording durations. Colored bars and black error bars indicate mean ± standard deviation. **J**: Computing time for parameter estimation as function of number of neurons. If not stated otherwise, parameter values were *N* = 10, *τ* = 250 ms, 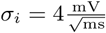 and *T* = 300 s.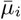, *C*_*i*_-pairs were drawn uniformly from the parameter space defined for Fig 1C. Here, *σ*_*i*_ was assumed to be known.

Classification performance (OUP vs. MJP) in terms of LLR for different values of population size, recording time, and process time constant is presented in Fig 2E. The dynamics of *x*(*t*) are correctly classified in all tests. An increased amount of observed data, either by population size or recording time, facilitates classification. Small values of the time constant for *x*(*t*) impede the decision, similarly as in the single neuron case (cf. Fig 1D, but note the difference in magnitude of the LLR).

Accuracy of parameter inference from multiple tests is shown in Fig 2F-I. The true values of 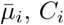, and *τ* are well approximated, and errors between true and estimated values increase only mildly as recording time is reduced from 10 min to 2.5 min.

Computation times for parameter inference are visualized in Fig 2J. From the tests considered in this section computation time appears to increase linearly with the number of neurons in the population due to the efficient optimization scheme that allows for parallelization over observed neurons.

We next examine whether our inference method serves to capture qualitatively distinct shared dynamics. For this purpose we consider synthetic data from simulations of the population model where *x*(*t*) is described by a sine wave instead of the OUP or MJP (Fig 3). The results from these examples demonstrate that the oscillatory dynamics of *x*(*t*) can be well recovered from a population of *N* = 20 neurons using our method based on either the OUP or MJP. Reconstruction accuracy deteriorates with increasing oscillation frequency; furthermore, the task becomes increasingly challenging for small populations and small neuronal spike rates. In sum, our method allows for accurate parameter inference, reconstruction and classification of the shared input dynamics, and spike rate estimation based on synthetic population data. It further serves to reconstruct oscillatory dynamics of the shared input, which demonstrates the dynamical flexibility of the doubly-stochastic I&F model.

**Figure 3.**
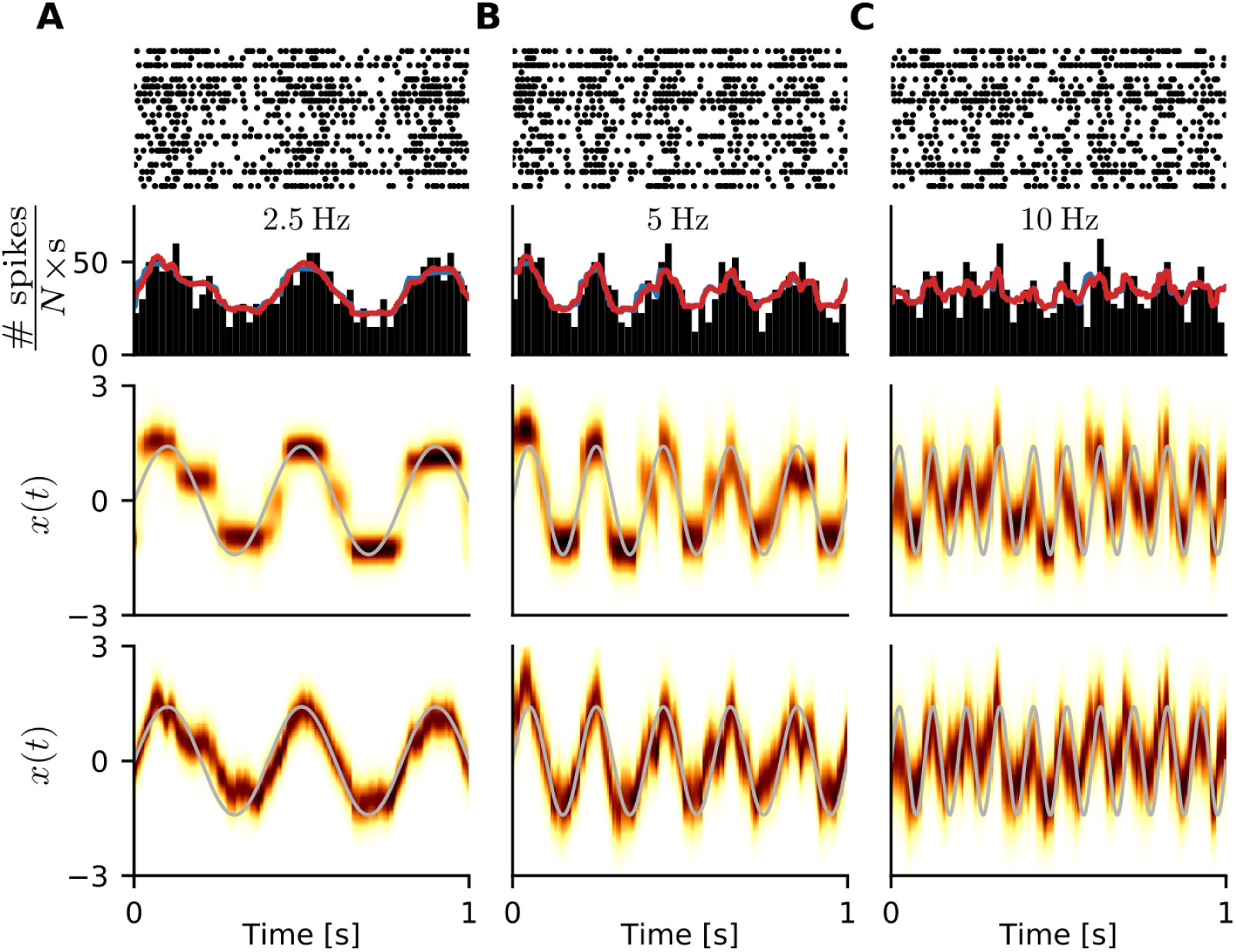
Inference results for neuronal populations driven by oscillating common input. From top to bottom: Spike trains of an I&F population with shared input component *x*(*t*) described by a sine wave, population spike rate histogram and estimates from our method using the OUP (red line) or MJP (blue line), true time series of *x*(*t*) (grey line) and sequence of the inferred density over *x* using the MJP or OUP (below). Oscillation frequency was 2.5 Hz (**A**), 5 Hz (**B**) and 10 Hz (**C**). The amplitude of *x*(*t*) was chosen such that the signal has unit standard deviation. The values for 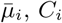, and *σ*_*i*_ were generated as in Fig 2, other parameter values were *N* = 20, *T* = 300 s.

### 4 Validation based on in-vitro ground truth recordings

We validate our method using ground truth data from patch-clamp recordings of cortical pyramidal neurons [27]. The cells received an injected current composed of a weak sinusoidal signal with strong additive noise such that neuronal spike rates oscillated roughly between 2 and 6 spikes/s (for details see Methods section 6). We fitted the doubly-stochastic I&F model using only the spike times and compared the inferred oscillatory dynamics of the mean input with the true oscillation of the current signal (Fig 4). The available data from three example recordings with different oscillation period and the corresponding results from our method are shown in Fig 4A-C. The magnitude of the rapid additive fluctuations (noise) dominates the oscillation amplitude of the injected current, so that spiking activity is only weakly modulated in an oscillatory manner. We measured the strength of this modulation by the resultant vector length (RVL) for the distribution of phases of the signal at spike times (Fig 4A-C right). The RVL quantifies the degree of concentration of that circular distribution; a large value of the RVL close to 1 indicates a strongly peaked, narrow distribution. Our inferred sequences of conditional density over the mean input clearly exhibit oscillatory dynamics. Importantly, the respective amplitude-normalized oscillations we extracted from these inferred sequences match well with the (amplitude-normalized) true signals (for details see Methods section 6).

**Figure 4.**
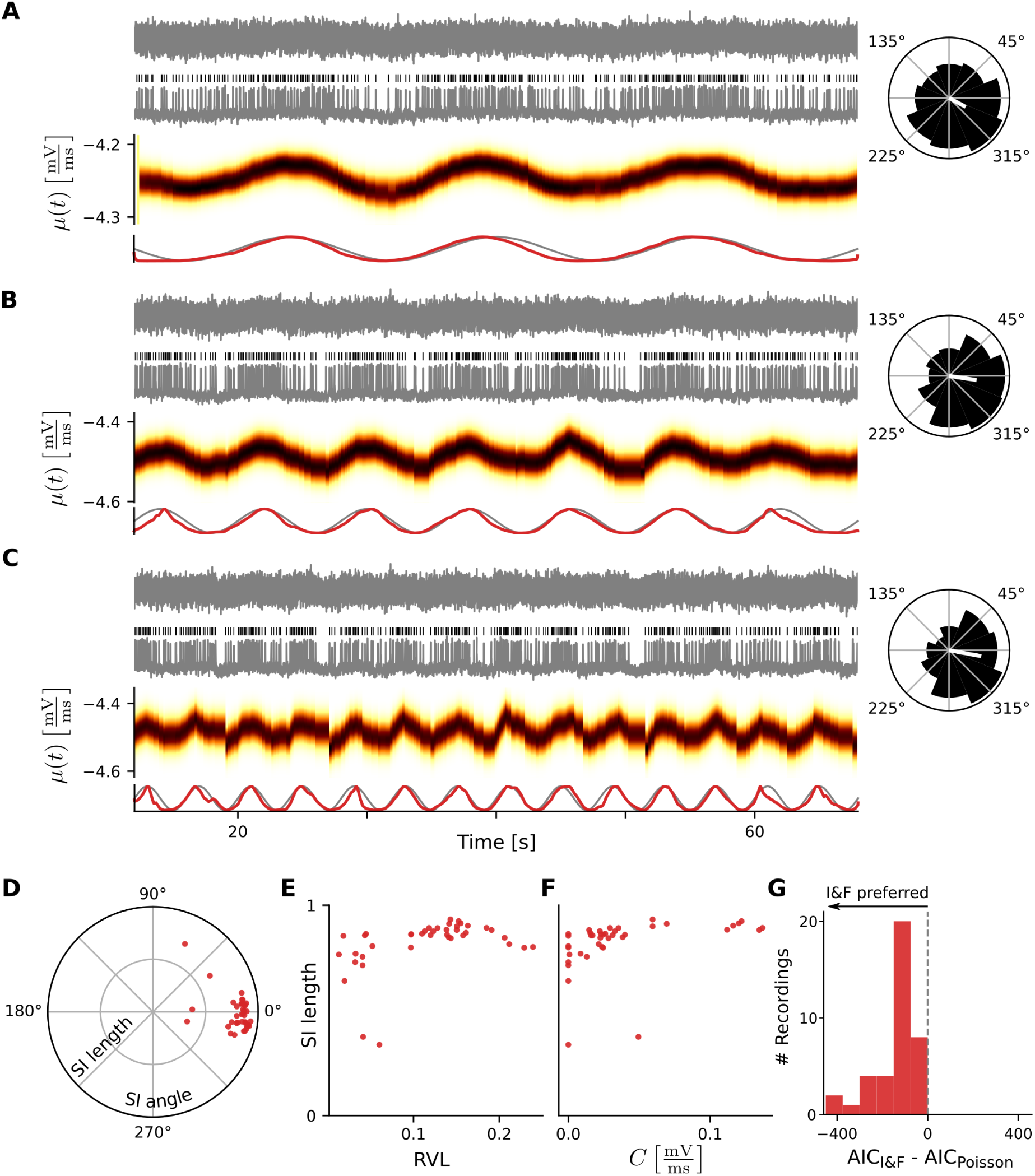
Inference results using in-vitro ground truth data. **A**–**C**, from top to bottom: Injected current, spike times and membrane voltage trace of a recorded neuron, evolution of the inferred density over the mean input, amplitude-normalized true (gray curve) and extracted (red curve) oscillatory signal (for details see Methods section 6). Inset: circular histogram of phases of the sinusoidal (true) input signal at spike times; the white line indicates the circular mean, its magnitude defines the resultant vector length (RVL). Oscillation period was 16 s (**A**), 8 s (**B**) and 4 s (**C**). **D** Synchrony index (SI) between inferred and true oscillatory signals, each dots corresponds to a recording. **E**: Synchronization strength |SI| (measuring the quality of signal reconstruction) versus RVL (measuring the locking strength between spikes and the oscillatory input signal). **F**: |SI| versus estimated value of coupling strength *C*. **G**: Difference between Akaike information criterion (AIC) for the I&F model and the Poisson model. In **D**–**G** all recordings with oscillation period of 16 s were used: 40 datasets from 13 different neurons. For inference the OUP was used to describe *x*(*t*) (also in the Poisson model).

We next quantified the synchronization between the extracted and true oscillatory signals using the synchrony index (SI), a complex number whose radius (magnitude) indicates the strength of locking and the angle represents the average phase shift between the oscillations (for details see Methods section 6). The SI for all cells is shown in Fig 4D. The angle values are concentrated around 0 and most magnitudes are large, close to 1, which means that the oscillations are aligned locking is strong for most cells. The locking strength, which measures the quality of signal reconstruction, clearly depends on the extent of oscillatory structure in the spiking activity, as quantified by the RVL (Fig 4E). Those few cells for which locking is weak exhibit small RVL values, for RVL values ≳ 0.1 signal reconstruction is excellent. Weak locking is further reflected by a small value of inferred coupling strength *C* (Fig 4F).

Finally, we compared the fitting performance of the I&F model with that of an inhomogeneous Poisson process using the AIC on these data (Fig 4G, cf. Results section 2). The preferred model is indicated by the lower AIC value. For all recordings the I&F model outperforms the Poisson model according to this measure.

In this section we used the OUP for the latent process *x*(*t*) (in the I&F and Poisson models); using instead the MJP did not significantly change the results. Note that the models do not include any prior information about oscillatory dynamics.

In sum, our method allows to accurately recover the ground truth dynamics of hidden, weak input signals from spike trains of cortical neurons and outperforms a classical model-based approach on these data.

### 5 Application to in-vivo multi-electrode recordings

We now apply our method to extracellular recordings from primary visual cortex of a macaque monkey under anaesthesia [28]. The data consist of spontaneous spiking activity of a population of *N* = 20 single units over a duration of 10 minutes, which was split into two sets of 5 minutes each (one for fitting and one for testing; for details on preprocessing see Methods section 7). We fitted both I&F model variants, with either OUP or MJP for the shared input dynamics, to these spike trains and compared their fitting performance on the test set. The MJP is favored according to the LLR evaluated on the test data (LLR_test_ = −443.0). Notably, a benchmark test against a Poisson point process model demonstrates a clear preference for the I&F population (LLR_test_ = −1514.9, using the MJP in both models). A 30 s segment of the spike-train data from the test set and the results from our method are shown in Fig 5A-C. The spike trains are ordered according to the inferred value of coupling strength *C*_*i*_ (Fig 5A). Units which strongly couple to the shared input component exhibit coordinated spiking activity whereas spiking of units with weak coupling appears more disordered. Using the inferred coupling strength *C*_*i*_, which quantifies the neuronal contribution to the low-dimensional population dynamics, we can thus separate “chorister” from “soloist” neurons [37] in a statistically principled way.

**Figure 5.**
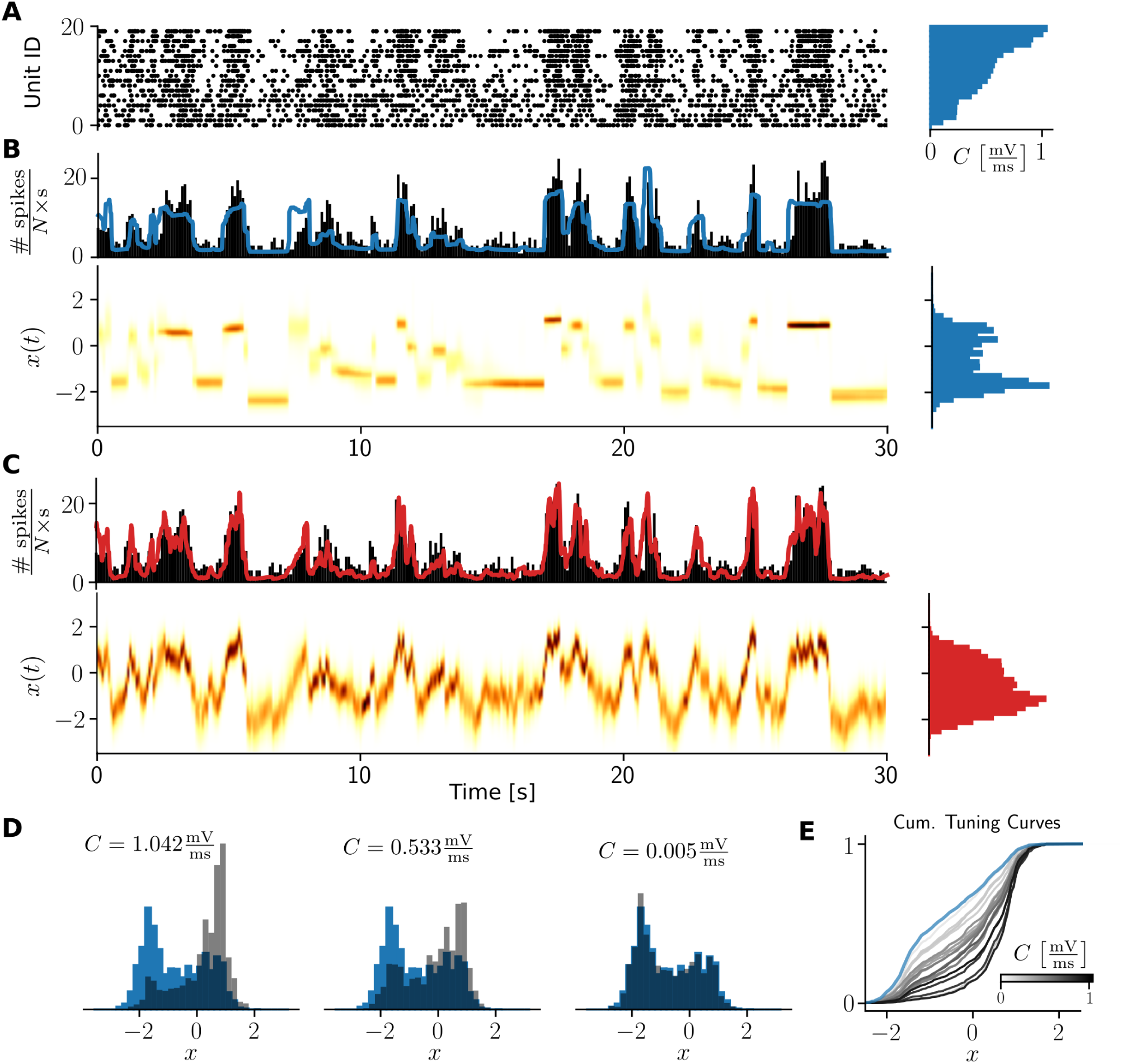
Inference results using in-vivo population data. **A**: Spike trains of 20 single units sorted according to the inferred coupling strength *C*_*i*_ (right). **B** and **C**, top: Population spike rate histogram and estimated instantaneous rate (colored line) using the MJP (**B**) or the OUP (**C**). Bottom: evolution of the inferred density over the common latent variable *x*. Right: Normalized histogram of expected values of *x* summed across the entire test set duration (latent state distribution). **D**: Normalized histogram of expected values of *x* at spike times (grey) with large (left), intermediate (center), and small estimated *C*_*i*_ (right). The latent state distribution is shown in blue (same as **B** right). **E**: Cumulative distribution functions corresponding to the histograms **D** for all single units, color-coded according to the value of estimated coupling strength. The Blue curve indicates the latent state distribution.

Population rate estimates versus empirical rate histograms are depicted in Fig 5B,C. Surprisingly, the OUP model variant yields a slightly better match in this respect despite its inferior fitting performance. Note, however, that approximating the population rate histogram is not an explicit objective of the fitting procedure.

Histograms over the sequence of expected values of *x* across the entire recording (which we refer to as latent state distributions) are shown in Fig 5B,C right. Note that the distributions differ from a standard normal, which is the stationary distribution of *x* as expected from the generative model. This indicates that the inferred low-dimensional dynamics deviate from both pure model variants for *x*, and it shows that our model flexibly captures different dynamics (as clearly demonstrated above, cf. Figs 3 and 4). Using only the values of the sequence of expected *x* at the observed spike times of a neuron we obtain *N* = 20 distinct histograms (one for each unit). This allows us to measure the extent to which individual neurons are tuned to the estimated shared dynamics in a way that does not directly depend on the inferred coupling strengths. Comparison between that measure of tuning and the coupling strengths enables another (indirect) validation. Indeed, units with a large value of *C* exhibit strong tuning, shown by distributions that are skewed towards large values of *x* (and clearly differ from the latent state distribution), while units with weak coupling also exhibit weak tuning, shown by distributions that are indistinguishable from the latent state distribution (Fig 5D,E).

In sum, our method allows to characterize the low-dimensional population dynamics and the contribution/coupling of individual neurons to those dynamics in a principled way.

## Discussion

We presented a statistically principled approach to fit an I&F population model with doubly-stochastic input to spike-train data. The input contains neuron-specific white noise fluctuations and a shared Markovian component that dominates the low-dimensional overall population dynamics. We extensively evaluated our methodology on synthetic data, validated it using ground truth in-vitro and in-vivo recordings, and compared the I&F population to a classical Poisson point process model in terms of fitting performance. Altogether, the results demonstrated the benefits of our approach to identify and classify the latent low-dimensional population dynamics, and quantify how individual neurons couple to those dynamics.

### Related Work

At the core of our methodology we exploit efficient numerical schemes to compute the ISI probability density of an I&F neuron exposed to Gaussian white noise input with high precision. The ability to rapidly compute this density allows to evaluate the likelihood of observed spike trains, which has been previously used for inference purposes using single stochastic I&F neurons [25, 38] and networks [25]. The nonstationarity considered here (doubly-stochastic population model) constitutes a relevant and highly nontrivial extension.

A related approach on the level of single neurons has been introduced in [33], which uses a diffusion process for the dynamics of the mean input and involves an approximation of the ISI probability density to avoid its numerical computation. An advantage of our approach in this regard is that it allows for the application of various, reasonable processes for the mean input, as demonstrated using the OUP and MJP. Note that the variance of these processes remains fixed over time, whereas that of a diffusion process increases without bounds. Importantly, we further considered neuronal populations.

Methodologically related work previously focused on the latent dynamics that underlie the spiking activity of single neurons, using Poisson point process models with continuous or jumping dynamics of the underlying rate, similar to those considered here [39, 40]. In our comparisons the doubly-stochastic I&F model clearly outperformed such Poisson point process models with latent rate dynamics. Whether and in what way this could affect classification results [40] is currently not clear.

Abstract models with larger complexity, in particular, generalized linear point process models that account for the dynamics of latent variables can be designed and optimized to fit well observed spike train data [9, 41–44]. However, the dynamical flexibility of these models is typically associated with large numbers of parameters that need to be optimized; hence, this approach is prone to overfitting unless strong contraints and/or parameter regularization are enforced [7, 36, 45–47]. An advantage of our approach in this regard is that the parameter space for optimization is comparably low-dimensional, which diminishes the risk of overfitting, without sacrificing essential aspects of neuronal spiking dynamics. Furthermore, the variables and parameters of I&F models can be interpreted in a straightforward way.

### Limitations & potential extensions

In our inference method the mean input is approximated by one value for each neuron and each ISI. This approximation leads to an “event-based” (spike-based) temporal binning of the inferred sequences, as was previously used for single neurons [33]. It implies that the latent dynamics cannot be captured precisely across extended intervals without spikes.

We aimed to capture the latent low-dimensional collective dynamics using a stochastic process for the variations of shared inputs. This process describes the dynamics of the external drive and picks up effects due to synaptic interactions on the population-average level. Detailed effects of synaptic coupling between the observed neurons are not explicitly included. The assumption that inputs from unobserved neurons dominate the collective population dynamics is well justified as long as the observed neurons make up only a small fraction of the local population (which is a typical scenario) [10]. Nevertheless, the model we considered could be extended to a doubly-stochastic I&F circuit with an explicit description of synaptic interactions between observed neurons. In particular, a combination of our approach with recent inference methods for I&F circuits [25] – for example, using the inferred (sequences of) parameters of our population model for subsequent estimation of connectivity – may be beneficial for the estimation of synaptic couplings.

We used the classical leaky I&F model to describe the dynamics of the neuronal membrane voltage (with focus on spiking). Our methodology, however, can be easily extended to other I&F model variants for which spike emission is a renewal process, such as the exponential I&F model [48]. Furthermore, an absolute refractory period can be included in a straightforward way.

We considered two qualitatively different Markov processes (OUP and MJP) for the shared input fluctuations. Other processes may be used instead: for example, a differentiable process whose temporal correlation-function is described by a Matérn kernel or periodic kernel (see, e.g., [49]).

Here we considered a one-dimensional process for the dynamics of shared input variations, which dominate the population dynamics. Depending on the size of the observed population and application it may be relevant to account for multi-dimensional latent dynamics; our framework can be extended accordingly in a similar way as for generalized linear models [9].

### Applicability

Using in-vitro ground truth data we have demonstrated that our approach accurately extracts the latent dynamics of input signals that are masked by strong noise, outperforming a more classical model-based method. These results support the validity of our method for application to in-vivo data that typically do not entail ground truth about the properties to be inferred. For such a scenario, using spike train data from extracellular recordings, we have shown that the inferred coupling strengths allow to distinguish between cells which clearly participate in coordinated population activity and those which spike rather independently. Such a characterization of “chorister” and “soloist” neurons has recently been shown to correlate with the underlying synaptic connectivity [37]. In future applications our method may serve, for example, to relate classification results (between model variants) and inferred parameter values across different experimental conditions.

## Materials and methods

### 1 Generative model

We consider a population of *N* leaky I&F neurons driven by fluctuating inputs. The dynamics of the membrane voltage *V*_*i*_ of neuron *i* (*i* ∈ {1, …, *N*}) are governed by the stochastic differential equation

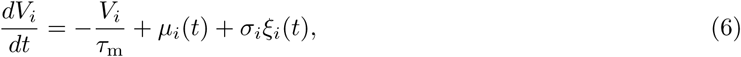

where *τ*_m_ denotes the membrane time constant. The input variations are described by the time-dependent mean *µ*_*i*_(*t*) and Gaussian white noise process *ξ*_*i*_(*t*) scaled by *σ*_*i*_, where ⟨*ξ*_*i*_(*t*)*ξ*_*j*_(*t* + Δ) ⟩ = *δ*_*i*,*j*_*δ*(Δ) for *i, j* ∈ {1, …, *N*} with expectation ⟨· ⟩. The neuron fires a spike when the membrane voltage reaches the spike threshold value *V*_s_, subsequently *V*_*i*_ is reset to the value *V*_r_.

The dynamics of the mean input *µ*_*i*_(*t*) are determined by a common process *x*(*t*) through

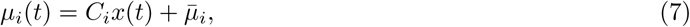

where *C*_*i*_ denotes the strength of coupling of neuron *i* to *x*(*t*) and 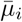 is the offset for neuron *i*. We separately consider two processes with qualitatively different dynamics for *x*(*t*): an *Ornstein-Uhlenbeck process* (OUP), described by

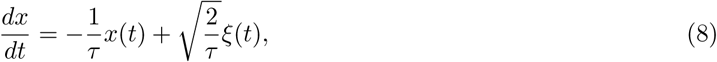

with time constant *τ* and Gaussian white noise process *ξ*(*t*), ⟨*ξ*(*t*)*ξ*(*t* + Δ) ⟩ = *δ*(Δ); alternatively, *x*(*t*) is described by a *Markov jump process* (MJP) which is piece-wise constant across intervals with exponentially distributed duration of mean *τ* and takes values drawn from a standard normal distribution *𝒩*(0, 1),

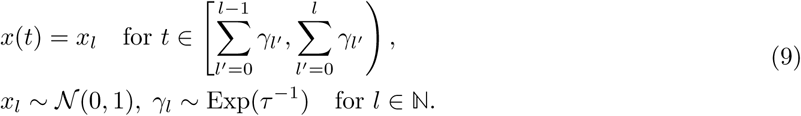

This process is also known as marked (homogeneous) Poisson point process. For both processes the stationary distribution is standard normal, lim_*t*→*∞*_ *x*(*t*) ∼*𝒩*(0, 1), and the autocorrelation function is given by

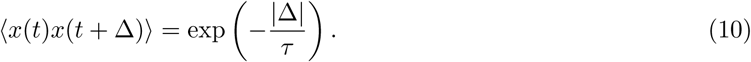

Hence, our doubly-stochastic model incorporates fast independent input fluctuations by *σ*_*i*_*ξ*_*i*_(*t*) in Eq (6) with Gaussian white noise process *ξ*_*i*_(*t*) and slower shared input variations by *µ*_*i*_(*t*) via Eq (7) with common OUP or MJP *x*(*t*).

It is not meaningful to estimate all parameters of this model. A change of *V*_s_ or *V*_r_ can be completely compensated in terms of spiking dynamics by appropriate changes of 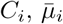 and *σ*_*i*_, which can be seen using the change of variables 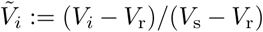. Consequently, we exclude *V*_s_ and *V*_r_ from the parameters to be inferred and instead set them to reasonable values. Furthermore, we fix *τ*_m_ for most of our results, since a change of that parameter can also be well compensated for by appropriate changes of the input parameters (see Results section 2). The parameter values were *τ*_m_ = 10 ms, *V*_s_ = −40 mV and *V*_r_ = −65 mV.

### 2 Statistical modeling

#### Spike-based discretization

We consider having observed the spike trains of *N* neurons and collect these data in a sequence of ISIs *s*_1:*K*_ := (*s*_1_, …, *s*_*K*_) that are ordered increasingly according to the last spike time of each ISI. Under the assumption that the process *x*(*t*) evolves slowly compared to the duration of the ISIs we approximate the mean input for each neuron across each ISI effectively by one value: 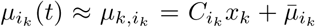 for all times *t* within the *k*-th ISI *s*_*k*_ in the sequence *s*_1:*K*_, where *x*_*k*_ is the value of *x*(*t*) at the end spike time of *s*_*k*_ and *i*_*k*_ indicates the neuron that corresponds to that ISI. Note that in order to simplify notation we use that *s*_*k*_, explicitly the duration of the *k*-th ISI, informs also (implicitly) about the start and end times of the ISI.

Hence, for the inference problem *x*(*t*) is replaced by the sequence *x*_0:*K*_ := (*x*_0_, …, *x*_*K*_) with *x*_0_ := *x*(0) and *x*_*k*_ := *x*(*t*_*k*_) for *k* ≥ 1, *t*_*k*_ being the *k*-th spike time across the population, which greatly facilitates the task.

#### Markov property

Due to the fact that both processes, OUP and MJP, are Markovian we can factorize the probability density of the sequence given the time constant parameter as

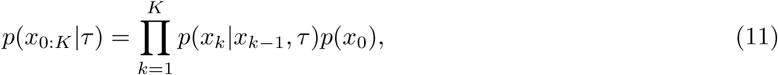

where *p*(*x*_0_) is the probability density for the initial process value (prior). We assume *p*(*x*_0_) ∼ *𝒩* (0, 1) which corresponds to the stationary distribution of the process. The transition probability density for the OUP is given by

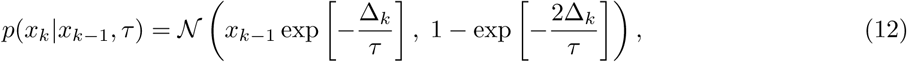

where Δ_*k*_ = *t*_*k*_ - *t*_*k-*1_. For notational convenience we do not explicitly indicate Δ_*k*_ in *p*(*x*_*k*_|*x*_*k-*1_, *τ*) and instead use that *x*_*k*_ also contains information about the corresponding time point. For the MJP, on the other hand, we have

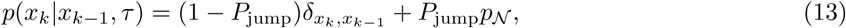

where the probability of jumping is 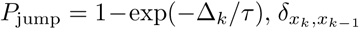 is the Kronecker delta and *p*_*𝒩*_ ∼ *𝒩*(0, 1). Note that *P*_jump_ denotes the probability that at least one jump occurs during an interval of duration Δ_*k*_.

#### Conditional likelihood

Since spike emission for an I&F neuron is a renewal process, the likelihood of the observed data *s*_1:*K*_ conditioned on the sequence *x*_0:*K*_ (which approximates the process *x*(*t*)) and neuron-specific parameter values can be factorized as

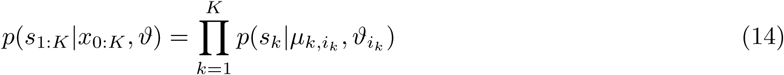

with 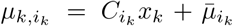. *ϑ* = {*ϑ*_1_, …, *ϑ*_*N*_} contains the parameters for each neuron, where 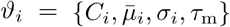. Each factor on the right hand side is the ISI probability density of an I&F neuron exposed to Gaussian white noise input with constant parameter values, evaluated at one point. We can compute the ISI density 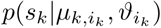 efficiently with high precision for a range of values *s* ≥ 0 at once. This is achieved by solving a Fokker-Planck partial differential equation that describes the so-called first passage time problem for the stochastic I&F model [25, 38, 50]. Here we apply the finite volume solution method described in [25]. In practice, we pre-compute 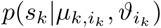 on a sufficiently fine grid of values for *s*_*k*_ and 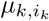, which is then used as a look-up table in the optimization procedure described below. For values of the mean input 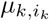 encountered during the inference procedure that are not on the grid we use linear interpolation. For additional details we refer to our Python implementation available at Github: https://github.com/neuromethods/inference_for_doubly_stochastic_IF_models

#### Joint and marginal likelihoods

By combining Eqs (11) and (14) we obtain the joint likelihood of the observed data and the process sequence as

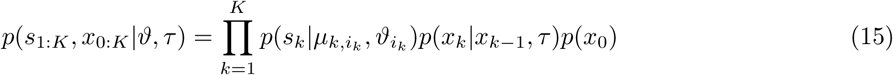

with 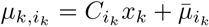. For the marginal likelihood of observing the data from the model, integration over *x*_0:*K*_ is required:

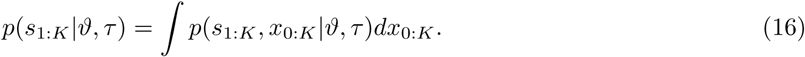

Note the difference between this marginal likelihood and the likelihood in Eq (14) which is conditioned on knowledge of the process sequence.

### 3 Parameter estimation

Estimates for a particular set of parameters are obtained by maximizing the marginal likelihood with respect to those parameters (maximum likelihood estimation). In the following we describe how we evaluate this likelihood from Eq (16) and how we maximize it with respect to the parameters of interest. For additional details we refer to our code available at Github.

#### Evaluation of the marginal likelihood

Integration over *x*_0:*K*_ in Eq (16) can be efficiently performed by an iterative procedure. Knowing *p*(*x*_*k-*1_|*s*_1:*k-*1_, *ϑ, τ*) allows for the prediction step using the Chapman-Kolmogorov (forward) equation

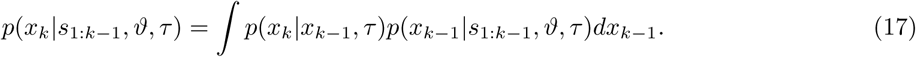

The iteration is initialized for *k* = 1 and *p*(*x*_0_|*s*_1:0_, *ϑ, τ*) = *p*(*x*_0_). In the following step we incorporate the next observation *s*_*k*_ by the filtering

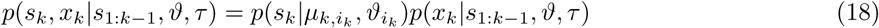

with 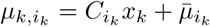. By marginalizing out *x*_*k*_ we obtain

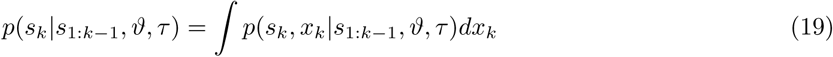

and calculate in the last step

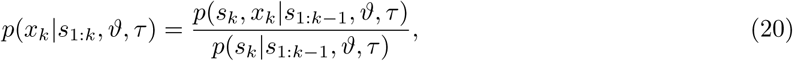

which completes the iteration *k* − 1 → *k*. After *K* iterations the marginal likelihood is given by

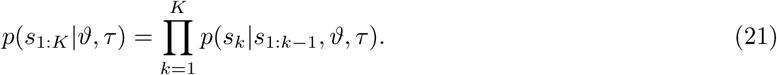

Practically, the steps of the iteration described above are performed using a reasonable discretization for *x*; here we used bins of width 0.05 in the range [−3.5, 3.5]. Consequently, the step in Eq (17), for example, involves a square matrix for the transition probability density and a sum for the integral.

#### Likelihood maximization for a single neuron

We maximize the logarithm of the likelihood from Eq (21) with respect to the parameters 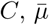 and *τ* using a simplex optimization method [35]. Note that our approach is not restricted to the simplex method, other (e.g. gradient-based) techniques may also be applied. The parameter *σ* is estimated via an outer loop, where we iterate over a range of values for *σ* and maximize the likelihood for each of those. The membrane time constant *τ*_m_ is not optimized as justified in Results section 2.

#### Likelihood maximization for a population

The parameter space for optimization has 3*N* + 1 dimensions, where *N* is the number of neurons. To reduce computational efforts and obtain parameter estimates in reasonable time we applied the following approximate optimization scheme. First, we estimate the noise amplitude *σ*_*i*_ for *i* ∈ {1, …, *N*} by fitting a doubly-stochastic I&F model with independent process *x*_*i*_(*t*) to each spike train separately, as described in the previous paragraph. Then, we re-estimate the offset 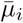 by fitting a model neuron with *C*_*i*_ = 0 and *σ*_*i*_ from the previous step for each neuron. Note that these two steps can be performed in parallel across neurons. Next, we use the full population model and maximize the likelihood with respect to all coupling strengths *C*_1:*N*_ := (*C*_1_, …, *C*_*N*_) given *τ* and vice versa in an alternating way, keeping *σ*_1:*N*_ and 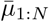 fixed. The optimization with respect to *C*_1:*N*_ is done in parallel across neurons: for optimizing *C*_*i*_ the couplings for other neurons are fixed to the values found in the previous optimization step. Finally, after convergence of the previous iteration procedure for *C*_1:*N*_ and *τ*, our estimates for the parameters *C*_1:*N*_ and 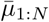 are improved by maximizing the likelihood with respect to *C*_*i*_ and 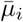 using the previously determined values as starting points, keeping all other parameters fixed, for *i* ∈{1, …, *N*}. This last step is performed again in parallel. Although this scheme appears rather approximate it yields accurate parameter estimates (cf. Results section 3).

### 4 Classification of latent dynamics and model comparison

To classify the dynamics of the slow (shared) input variations we compare both fitted model variants, one using the OUP and one using the MJP, by applying the log-likelihood ratio (LLR). This quantity can be expressed as the difference

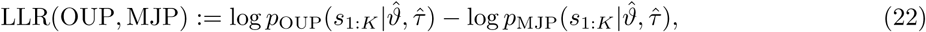

where the subscripts OUP and MJP indicate the considered process, and 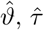 denote the parameter estimates determined prior to model comparison. Note that the LLR does not take into account the model complexity, which is not needed because both compared models have the same number of estimated parameters.

To compare models with different complexities in terms of number of parameters to be estimated, specifically the I&F and the Poisson models considered here, we apply the Akaike information criterion (AIC) [51]. This measure is given by 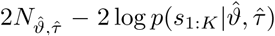, where 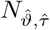 is the number of estimated parameters. The preferred model is indicated by the smallest AIC value.

### 5 Reconstruction of time series

To estimate the latent state sequence *x*_0:*K*_ we compute the probability density of the hidden process sequence given all observed ISIs and the model with inferred parameters 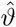 and 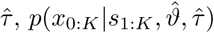. Specifically, we calculate the marginal densities conditioned on the observations and the fitted model, 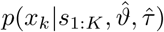 for *k* ∈ {1, …, *K*}, using

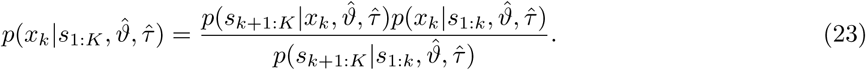

The probability density 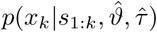 on the right hand side of Eq (23) is already known from the forward (filtering) iteration (cf. Eq (20)) at this point. We calculate the probability density 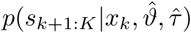 iteratively by backward smoothing,

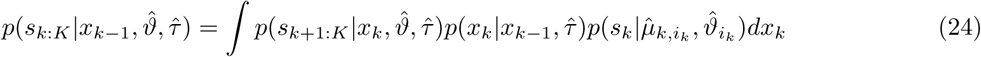

with 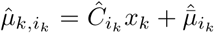. We initialize 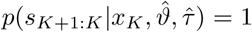 (uniform distribution), since *s*_*K*_ is the last observed ISI. The denominator in Eq (23) is obtained by numerical marginalization. In this way we obtain the sequence 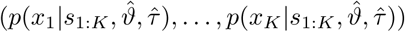 that we visualize in Figs 1–5. Using the elements from this sequence we estimate *x*_0:*K*_ by the sequence of expected values (⟨*x*_0_ ⟩, …, ⟨*x*_*K*_⟩), where *x*_0_ is calculated with respect to *p*(*x*_0_).

The sequence of marginal densities further allows us to estimate the (instantaneous) population spike rate time series by the sequence (*r*_0_, …, *r*_*K*_),

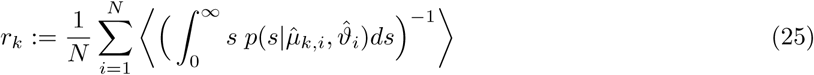

with 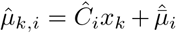, where the expectation ⟨·⟩ is calculated with respect to *p*(*x*_*k*_|*s*_1:*K*_, *ϑ, τ*). Each summand in Eq (25) is the expected inverse mean ISI a neuron at the time that corresponds to *x*_*k*_.

### 6 In-vitro data

For validation purposes we considered recorded activity of cortical pyramidal neurons from mouse brain slices [27]. Each recorded neuron was stimulated by a fluctuating current with

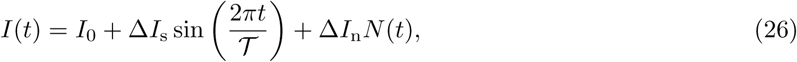

where *I*_0_ denotes the offset, Δ*I*_s_ the amplitude of the sinusoidal component with period 𝒯, and Δ*I*_n_ is the magnitude of the noise component *N* (*t*), which is an Ornstein-Uhlenbeck process with zero mean, unit variance and correlation time 3 ms. The parameters *I*_0_, Δ*I*_s_ and Δ*I*_n_ were tuned such that the neuronal spike rate oscillated between 2 and 6 spikes/s. We considered recordings that resulted in an average spike rate of at least 4 spikes/s. Oscillation period was 𝒯 ∈ {4, 8, 16} s. Each recording lasted 68 s: for the first 4 s the input current was constant, for the remaining 64 s it was described by Eq (26). For each of these 64 s segments we fitted a doubly-stochastic I&F model using the OUP variant. We aimed to capture the oscillatory dynamics by the OUP *x*(*t*) and approximated the fast fluctuations of the true input (note the small correlation time) by the Gaussian white noise process in the model. We determined the optimal noise amplitude *σ* in the model by maximizing the likelihood for a range of values of *σ* in the interval 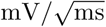.

To quantify oscillations in the inferred input and compare them with the true signal we first computed the time series of the expected mean input 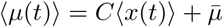 using the density 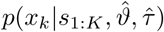 (cf. Methods section 5) and linear interpolation for values of the time series between spike times. Due to strong onset transients in the spiking activity we disregarded the first 8 s of the time series and linearly detrended ⟨*µ*(*t*)⟩ (using the function `signal.detrend` from the Python package `scipy`) for these comparisons.

The instantaneous phase of this time series, *ϕ*_est_(*t*), was then obtained using the Hilbert transform. Analogously, the instantaneous phase of the true signal, *ϕ*_true_(*t*), was calculated using the Hilbert transform of the sinusoidal input component. To visualize the oscillations in the true and inferred time series we used sin(*ϕ*_true_(*t*)) and sin(*ϕ*_est_(*t*)), respectively (Fig 4A-C). We next computed the synchrony index (SI) [52] as

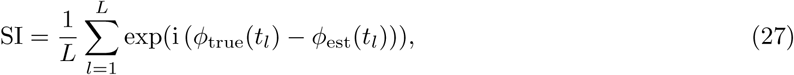

where *t*_1_, …, *t*_*L*_ are the time points of experimental measurements. The radius (or magnitude) of the SI measures the strength of locking and the angle indicates the average phase shift between the two oscillations.

To measure the strength of locking of neuronal spiking to the (true) oscillatory signal we calculated the resultant vector length (RVL) of the circular mean of phase at spike times *t*_0_, …, *t*_*K*_,

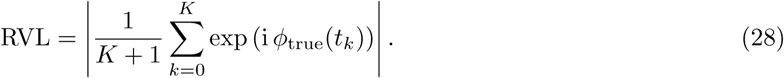

### 7 In-vivo data

We applied our method to extracellular recordings from primary visual cortex of an anaesthetized monkey [28, 53]. Extracellular potentials were recorded using an implanted multi-electrode array. After spike sorting, the data consisted of spike times of several single and multi-units. Following [53], we discarded units with signal-to-noise ratio < 2.75 (average wave form amplitude divided by the standard deviation of the wave form noise) as multi-units. We further excluded units with average spike rate > 100 Hz and omitted unphysiologically short ISIs < 3 ms. This resulted in 26 single units that we used for fitting. To account for potential spike sorting errors in this dataset we replaced the factors of the conditional likelihood in Eq (14) by 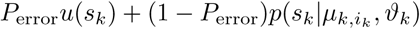, where *P*_error_ is the probability of a spike sorting error and *u*(*s*) is the density of a uniform distribution over a reasonably bounded interval. We set *P*_error_ = 0.05 in consistency with experimental observations [54]. We first fitted a doubly-stochastic I&F neuron for each of these units and determined the optimal value of *σ*_*i*_ using the OUP and MJP separately. For population analyses we then considered the 20 single units with the slowest inferred processes (i.e., those with the largest time constant *τ*, using averages across the results from the OUP and MJP model variants). We continued the fitting procedure for that population as described in Methods section 3.

## Acknowledgments

We thank Christian Pozzorini for making available the in-vitro data. CD was supported by the German Research Foundation via GRK1589/2 and CRC1295.

## Supporting information

**Figure S1.**
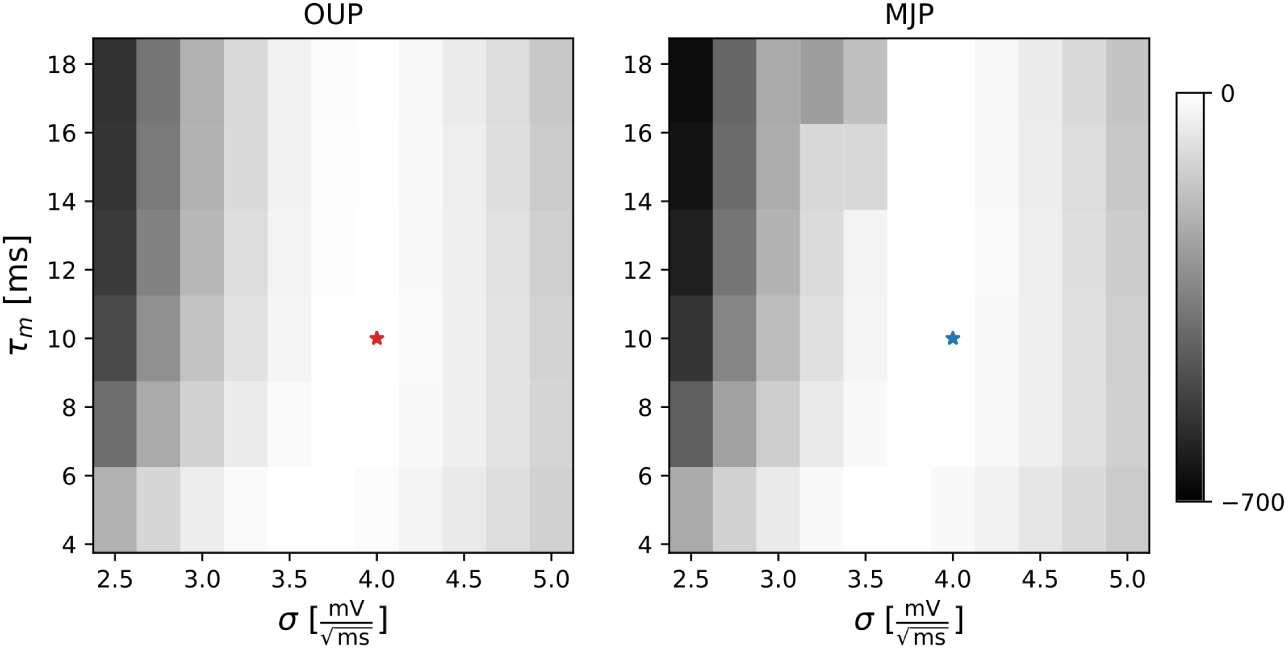
Dependence of maximal likelihood on membrane time constant and noise intensity. Difference between maximal log-likelihood for different values of *τ*_m_ and *σ*, and the one obtained for the true parameter values using the OUP (**A**) or the MJP (**B**). Colored stars mark the true parameter values of *τ*_m_ and *σ*. Other parameter values were 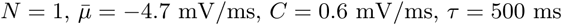, and 5 *×* 10^3^ spikes are in the data.

## Notes

https://github.com/neuromethods/inference_for_doubly_stochastic_IF_models

## References

1. Shadlen MN, Newsome WT. The Variable Discharge of Cortical Neurons: Implications for Connectivity, Computation, and Information Coding. J Neurosci. 1998;18(10):3870–3896.

2. Luczak A, Barthó P, Harris KD. Spontaneous Events Outline the Realm of Possible Sensory Responses in Neocortical Populations. Neuron. 2009;62(3):413–425. doi:10.1016/j.neuron.2009.03.014.

3. Churchland MM, Cunningham JP, Kaufman MT, Foster JD, Nuyujukian P, Ryu SI, et al. Neural population dynamics during reaching. Nature. 2012;487:51–56. doi:10.1038/nature11129.

4. Mante V, Sussillo D, Shenoy KV, Newsome WT. Context-dependent computation by recurrent dynamics in prefrontal cortex. Nature. 2013;503(7474):78–84. doi:10.1038/nature12742.

5. Gallego JA, Perich MG, Miller LE, Solla SA. Neural Manifolds for the Control of Movement. Neuron. 2017;94(5):978–984. doi:10.1016/j.neuron.2017.05.025.

6. Cunningham JP, Yu BM. Dimensionality reduction for large-scale neural recordings. Nat Neurosci. 2014;17(11):1500–1509. doi:10.1038/nn.3776.

7. Aljadeff J, Lansdell BJ, Fairhall AL, Kleinfeld D. Analysis of Neuronal Spike Trains, Deconstructed. Neuron. 2016;91(2):221–259. doi:10.1016/j.neuron.2016.05.039.

8. Yu BM, Cunningham JP, Santhanam G, Ryu SI, Shenoy KV, Sahani M. Gaussian-Process Factor Analysis for Low-Dimensional Single-Trial Analysis of Neural Population Activity. J Neurophysiol. 2009;102(1):614–635. doi:10.1152/jn.90941.2008.

9. Lawhern V, Wu W, Hatsopoulos N, Paninski L. Population decoding of motor cortical activity using a generalized linear model with hidden states. J Neurosci Meth. 2010;189(2):267–280. doi:10.1016/j.jneumeth.2010.03.024.

10. Macke JH, Büsing L, Cunningham JP, Yu BM, Shenoy K, Sahani M. Empirical models of spiking in neural populations. Advances in Neural Information Processing Systems. 2011; p. 1350–1358.

11. Shimazaki H, Amari Si, Brown EN, Grün S. State-space analysis of time-varying higher-order spike correlation for multiple neural spike train data. PLoS Computational Biology. 2012;8(3). doi:10.1371/journal.pcbi.1002385.

12. Donner C, Obermayer K, Shimazaki H. Approximate Inference for Time-Varying Interactions and Macroscopic Dynamics of Neural Populations. PLoS Computational Biology. 2017;13(1):e1005309. doi:10.1371/journal.pcbi.1005309.

13. Zhao Y, Park IM. Variational Latent Gaussian Process for Recovering Single-Trial Dynamics from Population Spike Trains. Neural Computation. y2018; doi:10.1162/NECO.

14. Pandarinath C, O’Shea DJ, Collins J, Jozefowicz R, Stavisky SD, Kao JC, et al. Inferring single-trial neural population dynamics using sequential auto-encoders. Nature Methods. 2018;15(October):805–815. doi:10.1101/152884.

15. Duncker L, Bohner G, Boussard J, Sahani M. Learning interpretable continuous-time models of latent stochastic dynamical systems. arXiv preprint. 2019;.

16. Hodgkin AL, Huxley AF. A quantitative description of membrane current and its application to conduction and excitation in nerve. Journal of Physiology. 1952;117(4):500–44.

17. Teeter C, Iyer R, Menon V, Gouwens N, Feng D, Berg J, et al. Generalized leaky integrate-and-fire models classify multiple neuron types. Nature Communications. 2018;9(1):709. doi:10.1038/s41467-017-02717-4.

18. Prinz AA, Billimoria CP, Marder E. Alternative to Hand-Tuning Conductance-Based Models: Construction and Analysis of Databases of Model Neurons. J Neurophysiol. 2003;90(6):3998–4015. doi:10.1152/jn.00641.2003.

19. Marder E, Taylor AL. Multiple models to capture the variability in biological neurons and networks. Nat Neurosci. 2011;14(2):133–138. doi:10.1038/nn.2735.

20. Jolivet R, Kobayashi R, Rauch A, Naud R, Shinomoto S, Gerstner W. A benchmark test for a quantitative assessment of simple neuron models. J Neurosci Meth. 2008;169(2):417–424. doi:10.1016/j.jneumeth.2007.11.006.

21. Badel L, Lefort S, Brette R, Petersen CCH, Gerstner W, Richardson MJE. Dynamic I-V Curves Are Reliable Predictors of Naturalistic Pyramidal-Neuron Voltage Traces. J Neurophysiol. 2007;99(2):656– 666. doi:10.1152/jn.01107.2007.

22. Gerstner W, Naud R. How good are neuron models? Science. 2009;326(5951):379–380. doi:10.1126/science.1181936.

23. Harrison PM, Badel L, Wall MJ, Richardson MJE. Experimentally Verified Parameter Sets for Modelling Heterogeneous Neocortical Pyramidal-Cell Populations. PLoS Computational Biology. 2015;11(8):e1004165. doi:10.1371/journal.pcbi.1004165.

24. Pozzorini C, Mensi S, Hagens O, Naud R, Koch C, Gerstner W. Automated High-Throughput Characterization of Single Neurons by Means of Simplified Spiking Models. PLoS Computational Biology. 2015;11(6):e1004275. doi:10.1371/journal.pcbi.1004275.

25. Ladenbauer J, McKenzie S, English DF, Hagens O, Ostojic S. Inferring and validating mechanistic models of neural microcircuits based on spike-train data. bioRxiv. 2018; p. 1–31. doi:10.1101/261016.

26. Rabiner LR. A Tutorial on Hidden Markov Models and Selected Applications in Speech Recognition. Proceedings of the IEEE. 1989;77(2):257–286. doi:10.1109/5.18626.

27. Pozzorini C, Naud R, Mensi S, Gerstner W. Temporal whitening by power-law adaptation in neocortical neurons. Nat Neurosci. 2013;16:942–948. doi:10.1038/nn.3431.

28. Kohn A, Smith MA. Utah array extracellular recordings of spontaneous and visually evoked activity from anesthetized macaque primary visual cortex (V1). CRCNS. 2016; doi:http://dx.doi.org/10.6080/K0NC5Z4X.

29. Einevoll GT, Franke F, Hagen E, Pouzat C, Harris KD. Towards reliable spike-train recordings from thousands of neurons with multielectrodes. Current Opinion in Neurobiology. 2012;22(1):11–17. doi:10.1016/j.conb.2011.10.001.

30. Brunel N, Van Rossum MCW. Lapicque’s 1907 paper: From frogs to integrate-and-fire. Biol Cybern. 2007;97(5-6):337–339. doi:10.1007/s00422-007-0190-0.

31. Brunel N. Dynamics of Sparsely Connected Networks of Excitatory and Inhibitory Spiking Neurons. J Comput Neurosci. 2000;8(3):183–208. doi:10.1023/A:1008925309027.

32. Mattia M, Del Giudice P. Population dynamics of interacting spiking neurons. Phys Rev E. 2002;66(5):19. doi:10.1103/PhysRevE.66.051917.

33. Kim H, Shinomoto S. Estimating nonstationary input signals from a single neuronal spike train. Phys Rev E. 2012;86(5):1–12. doi:10.1103/PhysRevE.86.051903.

34. Augustin M, Ladenbauer J, Baumann F, Obermayer K. Low-dimensional spike rate models derived from networks of adaptive integrate-and-fire neurons: Comparison and implementation. PLoS Computational Biology. 2017;13(6):e1005545. doi:10.1371/journal.pcbi.1005545.

35. Nelder JA, Mead R. A Simplex Method for Function Minimization. The Computer Journal. 1965;7(4):308–313. doi:10.1093/comjnl/7.4.308.

36. Pillow JW, Shlens J, Paninski L, Sher A, Litke AM, Chichilnisky EJ, et al. Spatio-temporal correlations and visual signalling in a complete neuronal population. Nature. 2008;454(7207):995–999.

37. Okun M, Steinmetz NA, Cossell L, Iacaruso MF, Ko H, Barthò P, et al. Diverse coupling of neurons to populations in sensory cortex. Nature. 2015;521(7553):511–515. doi:10.1038/nature14273.

38. Mullowney P, Iyengar S. Parameter estimation for a leaky integrate-and-fire neuronal model from ISI data. J Comput Neurosci. 2008;24(2):179–194. doi:10.1007/s10827-007-0047-5.

39. Mochizuki Y, Shinomoto S. Analog and digital codes in the brain. Phys Rev E. 2014;89(2):022705. doi:10.1103/PhysRevE.89.022705.

40. Latimer KW, Yates JL, Meister MLR, Huk AC, Pillow JW. Single-trial spike trains in parietal cortex reveal discrete steps during decision-making. Science. 2015;349(6244):184–7. doi:10.1126/science.aaa4056.

41. Truccolo W, Eden UT, Fellows MR, Donoghue JP, Brown EN. A Point Process Framework for Relating Neural Spiking Activity to Spiking History, Neural Ensemble, and Extrinsic Covariate Effects. J Neurophysiol. 2005;93(2):1074–1089. doi:10.1152/jn.00697.2004.

42. Escola S, Fontanini A, Katz D, Paninski L. Hidden Markov Models for the Stimulus-Response Relationships of Multistate Neural Systems. Neural Computation. 2011;23(5):1071–1132. doi:10.1162/NECO_a_00118.

43. Vidne M, Ahmadian Y, Shlens J, Pillow JW, Kulkarni J, Litke AM, et al. Modeling the impact of common noise inputs on the network activity of retinal ganglion cells. J Comput Neurosci. 2012;33(1):97– 121. doi:10.1007/s10827-011-0376-2.

44. Tyrcha J, Roudi Y, Marsili M, Hertz J. The effect of nonstationarity on models inferred from neural data. Journal of Statistical Mechanics: Theory and Experiment. 2013;2013(3):P03005. doi:10.1088/1742-5468/2013/03/P03005.

45. Stevenson IH, London BM, Oby ER, Sachs NA, Reimer J, Englitz B, et al. Functional Connectivity and Tuning Curves in Populations of Simultaneously Recorded Neurons. PLoS Computational Biology. 2012;8(11):e1002775. doi:10.1371/journal.pcbi.1002775.

46. Zaytsev YV, Morrison A, Deger M. Reconstruction of recurrent synaptic connectivity of thousands of neurons from simulated spiking activity. J Comput Neurosci. 2015;39(1):77–103. doi:10.1007/s10827-015-0565-5.

47. Gerhard F, Deger M, Truccolo W. On the stability and dynamics of stochastic spiking neuron models: Nonlinear Hawkes process and point process GLMs. PLoS Computational Biology. 2017;13(2):e1005390. doi:10.1371/journal.pcbi.1005390.

48. Fourcaud-Trocmé N, Hansel D, van Vreeswijk C, Brunel N. How spike generation mechanisms determine the neuronal response to fluctuating inputs. J Neurosci. 2003;23(37):11628–11640.

49. Solin A. Stochastic differential equation methods for spatio-temporal Gaussian process regression. Aalto University. PhD Thesis; 2016. Available from: https://aaltodoc.aalto.fi/handle/123456789/19842.

50. Richardson MJE. Spike-train spectra and network response functions for non-linear integrate-and-fire neurons. Biol Cybern. 2008;99(4-5):381–92. doi:10.1007/s00422-008-0244-y.

51. Akaike H. A New Look at the Statistical Model Identification. IEEE Transactions on Automatic Control. 1974;19(6):716–723. doi:10.1109/TAC.1974.1100705.

52. Cohen MX. Assessing transient cross-frequency coupling in EEG data. J Neurosci Meth. 2008;168(2):494– 499. doi:10.1016/j.jneumeth.2007.10.012.

53. Smith MA, Kohn A. Spatial and Temporal Scales of Neuronal Correlation in Primary Visual Cortex. J Neurosci. 2008;28(48):12591–12603. doi:10.1523/jneurosci.2929-08.2008.

54. Rossant C, Kadir SN, Goodman DFM, Schulman J, Hunter MLD, Saleem AB, et al. Spike sorting for large, dense electrode arrays. Nat Neurosci. 2016;19(4):634–641. doi:10.1038/nn.4268.

